# Genome-Wide Association Analysis Identifies Genetic Correlates of Immune Infiltrates in Solid Tumors

**DOI:** 10.1101/106039

**Authors:** Nathan O. Siemers, James L Holloway, Han Chang, Scott D. Chasalow, Petra B. Ross-MacDonald, Charles F. Voliva, Joseph D. Szustakowski

## Abstract

Therapeutic options for the treatment of an increasing variety of cancers have been expanded by the introduction of a new class of drugs, commonly referred to as checkpoint blocking agents, that target the host immune system to positively modulate anti-tumor immune response. Although efficacy of these agents has been linked to a pre-existing level of tumor immune infiltrate, it remains unclear why some patients exhibit deep and durable responses to these agents therapy while others do not benefit. To examine the influence of tumor genetics on tumor immune state, we interrogated the relationship between somatic mutation and copy number alteration with infiltration levels of 7 immune cell types across 40 tumor cohorts in The Cancer Genome Atlas. Levels of cytotoxic T, regulatory T, total T, natural killer, and B cells, as well as monocytes and M2 macrophages, were estimated using a novel set of transcriptional signatures that were designed to resist interference from the cellular heterogeneity of tumors. Tumor mutational load and estimates of tumor purity were included in our association models to adjust for biases in multi-modal genomic data. Copy number alterations, mutations summarized at the gene level, and position-specific mutations were evaluated for association with tumor immune infiltration. We observed a strong relationship between copy number loss of a large region of chromosome 9p and decreased lymphocyte estimates in melanoma, pancreatic, and head/neck cancers. Mutations in the oncogenes PIK3CA, FGFR3, and RAS/RAF family members, as well as the tumor supressor TP53, were linked to changes in immune infiltration, usually in restricted tumor types. Associations of specific WNT/beta-catenin pathway genetic changes with immune state were limited, but we noted a link between 9p loss and the expression of the WNT receptor FZD3, suggesting that there are interactions between 9p alteration and WNT pathways. Finally, two different cell death regulators, CASP8 and DIDO1, were often mutated in head/neck tumors that had higher lymphocyte infiltrates. In summary, our study supports the relevance of tumor genetics to questions of efficacy and resistance in checkpoint blockade therapies. It also highlights the need to assess genome-wide influences during exploration of any specific tumor pathway hypothesized to be relevant to therapeutic response. Some of the observed genetic links to immune state, like 9p loss, may influence response to cancer immune therapies. Others, like mutations in cell death pathways, may help guide combination therapeutic approaches.

## Introduction

Checkpoint blocking cancer therapeutics, such as ipilimumab, nivolumab, pembrolizumab and atezolizumab, act by targeting immune cell signaling molecules rather than targeting the tumor directly. The molecular targets of these agents, CTLA-4, PD-1, and PD-L1, are components of pathways that inhibit T cell function[1]. Clinical experience with checkpoint blockade monotherapy and combinations has demonstrated dramatic tumor shrinkage and long-term durable, often drug-free, survival in some patients; however, many patients do not appear to benefit[2,3]. A number of different parameters have been explored to predict and explain the heterogeneity of patient benefit, within and across different cancer types. These include differences in the activation state of the tumor-immune infiltrate[4], differences in antigencity of the cancer cells due to differential expression and presentation of neo-antigens[5–8], and differences in composition of intestinal flora[9,10]. One of the most extensively studied potential biomarkers for checkpoint blocking agents is the cell surface expression of PD-L1, which is induced by interferon gamma from infiltrating lymphocytes and may be a surrogate for inflammatory state[11,12].

In addition to passenger mutations which can lead to expression of neo-antigens, the genetic history of tumorigenesis, manifest in the pattern of driver mutations and other necessary changes acquired during development, may affect the inflammatory state. Tumor driver pathways, such as WNT/Beta-catenin and FAK, have been recently linked with immune state in human tumors and identified as specific modulators of immune function in animal tumor models[13,14]. However, these studies have focused on specific cancer driver pathway hypotheses, and have yet to report their results in the context of a systematic genetic analysis. Rooney *et. al*. have reported a landmark study describing many tumor parameters, including genetic, that influence the strength of a ‘cytotoxic T cell signature’ in tumors across The Cancer Genome Atlas (TCGA, TCGA Research Network: http://cancergenome.nih.gov/)[15]. Porta-Pardo and Godzik have also studied the association of cancer mutations with a general estimate of immune infiltrate across TCGA[16]. Mutations associated with the interferon gamma signaling and antigen presentation pathways have recently been associated with acquired resistance to PD-1 blockade[8]. We sought to discern whether tumor genetic profiles could generally be correlated with the composition of specific subtypes of tumor immune infiltrate.

We have performed a systematic interrogation of the complex associations of cancer mutation and copy number alterations (CNA) with levels of immune infiltrate across solid TCGA tumors. Immune cell levels were estimated using novel transcriptional marker sets for major immune cell types, which were trained to resist interference from the cellular and transcriptional complexity of tumors. We also attempted to address several of the biases that introduce complexity in multi-modal TCGA data analysis in our analytical models. We have identified both previously reported and novel mechanisms by which tumor genetics may influence immune state. Additionally, we observed changes that suggest some tumors may genetically adapt to ongoing immune responses in ways that may broaden our definitions of immuno-editing[17]. Other associations of genetic changes with immune dynamics may reflect the evolutionary history of tumor subtypes in indirect ways, serving as a proxy for the tumors’ latent characteristics.

## Results

In our experience, many transcriptional marker sets for immune cells derived from the literature or trained from data on cell-sorted peripheral blood immune cells perform poorly when applied to data derived from tumor samples. Sometimes, even markers viewed as canonical for immune cell types are not always specific in RNA-seq data. For example, data from sorted human immune populations within the Immunological Genome Project suggests that CD4 transcription is higher on monocytes than on CD4+ T cells in RNA-seq data (http://immgen.org)[18,19]. Analysis of human melanoma single cell RNA-seq data from Tirosh et al. confirms that, in human tumors, CD4 is co-expressed with both T cell (CD3E) and myeloid (CSF1R) markers [20](Fig 1A).

**Figure 1.**
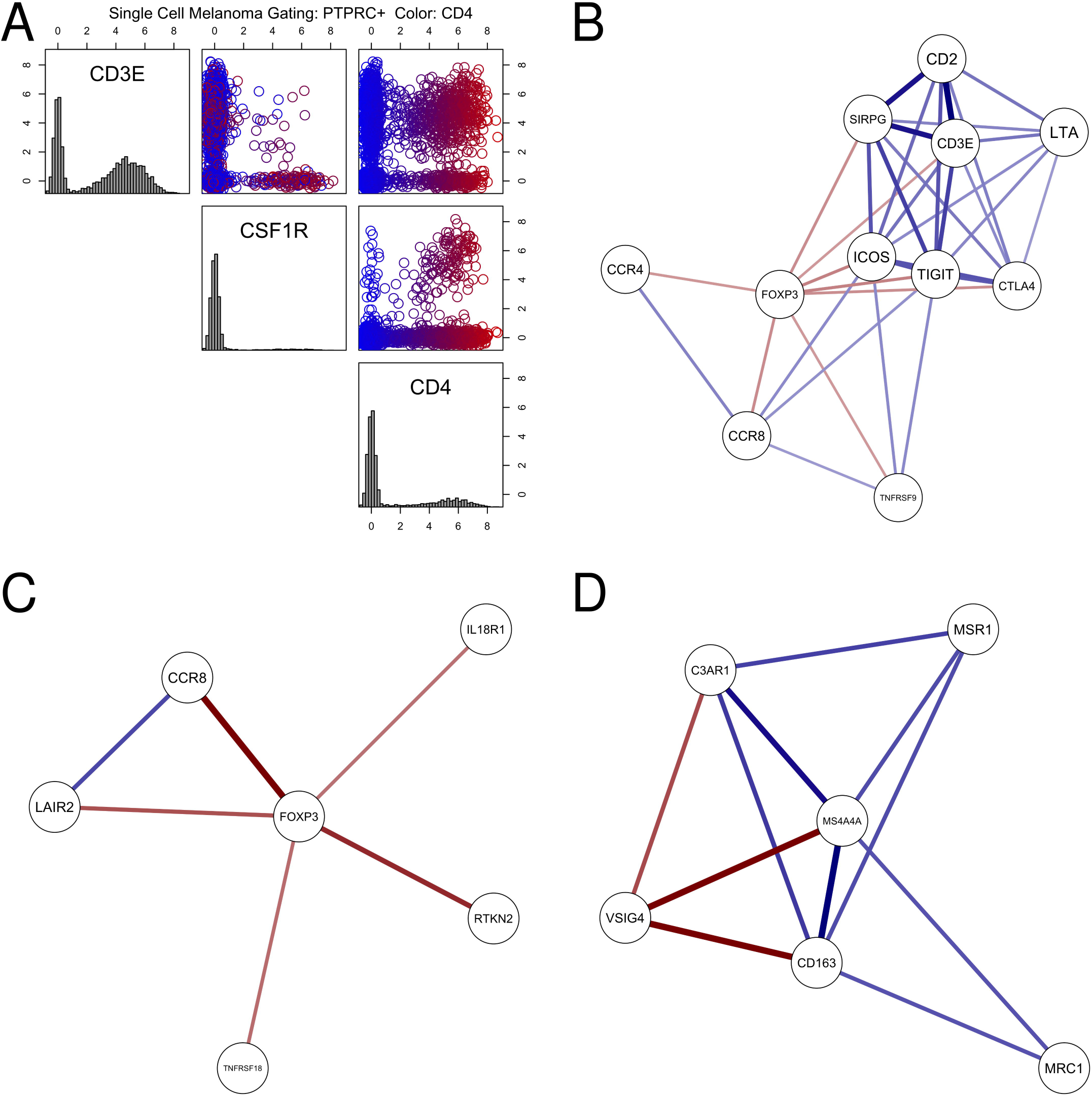
Expression and correlation of immunological markers in TCGA and Tirosh et al. single cell melanoma RNA-seq data. A: CD4 is co-expressed with both T cell (CD3E) and myeloid linege (CSF1R) markers in melanoma. Scatter plots of CD4, CD3E, and CSF1R transcript levels from a single-cell RNA-seq data study of melanoma patients (Tirosh et al.). Only CD45 positive cells (PTPRC, expression > 1) are shown. Gaussian noise (s.d. = 0.25) was added to the transcript estimates to improve data visualization (log2 scale). B: Mutual rank-based co-regulatory network around FOXP3 in TCGA. All solid tumor samples in the TCGA pan-cancer data release were used to create the mutual rank correlation network. Color saturation and thickness of lines represent strength of correlation. CCR8 and FOXP3 were selected to create a regulatory T cell (Treg) signature for estimating Treg content in tumors. C: Mutual rank-based co-regulatory network around FOXP3 in Tirosh et al. single cell melanoma RNA-seq data. D: Mutual rank-based co-regulatory network around macrophage marker VSIG4 in TCGA. VSIG4, CD163, and MS4A4A were selected to create a signature to estimate macrophage content in tumors.

In principle, if members of an immune marker set are cell-type specific, their transcript levels should be correlated with each other when examining a panel of tumors. We evaluated RNA expression correlation across TCGA solid-tumor cohorts for various immune cell marker sets obtained from the literature and from RNA-seq of sorted immune cells[21]. Our analysis often revealed very low correlation for expression of the genes within each given immune marker set, indicating that transcripts with coordinated expression in sorted immune cells are often heterogeneously expressed in more complex tissue samples. For example, a regulatory T cell (Treg) signature utilized in Rooney et al. was composed of seven genes, including the FOXP3 transcription factor, whose expression is a hallmark of CD4+ regulatory T (Treg) cells[15,22]. Pearson correlation analysis of the other members of this signature with FOXP3 across TCGA tumors revealed only 1 marker with correlation above 0.5 (CTLA4, 0.76) and two markers with correlation below 0.1 (IL4, 0.05; IL5, 0.01).

We therefore created alternative sets of transcriptional markers for immune cell types, designed to be appropriate to study of complex tumor samples. We did this by demanding that members of a set retain correlation with each other when examining TCGA tumors, and also that they are highly ranking neighbors of each other when looking at correlations of all transcripts in TCGA RNA-seq data. These sets were derived by first evaluating co-expression of candidate sentinel markers that displayed selectivity of RNA expression for the target cell type (CD8A for CD8+ T cells, FOXP3 for regulatory T cells, etc) with all transcripts across TCGA solid-tumor cohorts (details in Methods). We then used a stringent metric of mutual rank distance to identify gene neighbors for the sentinel markers[23] (Table 1). The principle of the mutual rank metric is that that highest scoring gene neighbors are not only highly correlated with each other, but are also each other’s highest ranking match, and a penalty is applied as as these mutual ranks become lower. These methods were employed to limit the inclusion of transcripts that are present in a more diverse range of cell types than the sentinel (manuscript in preparation).

**Table 1.** RNA-seq based marker sets created and used in this study to estimate immune cell levels in tumors.

| Signature | Membership | Description |
| --- | --- | --- |
| TCD8 | CD8A, CD8B | CD8+ T cell |
| Treg | FOXP3, CCR8 | Regulatory T cell |
| Tcell | CD3D, CD3E, CD2 | T cell (general) |
| Bcell | CD19, CD79A, MS4A1 | B cell |
| NK | KIR2DL1, KIR2DL3, KIR2DL4, KIR3DL1, KIR3DL2, KIR3DL3, KIR2DS4 | Natural killer cell |
| Mono | CD86, CSF1R, C3AR1 | Monocyte |
| MFm2 | CD163, VSIG4, MS4A4A | M2 Macrophage |
| TregCD8 | Treg, TCD8 | Treg versus CD8+ T cell |
| NKCD8 | NK, TCD8 | NK versus CD8+ T cell |

Figure 1B presents one example, a view of the (mutual rank) co-regulatory network around around FOXP3, a canonical marker of immuno-suppressive regulatory T cells. Transcript abundances for CCR8 possessed a both a strong as well as selective correlation to FOXP3 when compared to other neighbors, including transcripts probably more reflective of pan-T cell content (CD3 epsilon/CD3E, CD2), or markers correlated with both FOXP3 and pan-T markers (TIGIT, ICOS). A similar mutual rank analysis of the Tiros et al. single cell melanoma RNA-seq confirmed the co-expression of FOXP3 and CCR8 (Figure 1C). FOXP3 and CCR8 were ultimately chosen as a signature set for Treg estimation. Signature sets for other immune cell types were derived in a similar fashion; for macrophages, a highly co-regulated set (VSIG4, CD163, MS4A4A) was derived (Macrophage: Figure 1D). To create a quantitative score from the signature genes, simple medians of each marker set were used, once individual marker expression was normalized to a standard distribution (details in Methods).

We observed mutations, focal CNA, and large scale CNA, some spanning hundreds of millions of bases, associated with changes in estimated levels of immune cell types in tumors.For example, in TCGA head and neck cancer, we observed relationships between CD8+ T cell estimates and mutations in the p53 tumor suppressor (TP53, Chr17), caspase 8 (CASP8, Chr2) the RNA plymerase component POLR3A (Chr10), death-inducer obliterator 1 (DIDO1 Chr20), and CYLD (Chr16), a modulator of the nuclear factor kappa-B pathway (Fig 2). CD8 T cell levels were also in association with large copy number alterations on chromosomes 3, 5, 9, 18, and other chromosomal regions. Copy number gains along a region of chromosme 16 are in association with lower CD8 T cell levels, while copy number losses are associated with higher CD8 estimates. the peaks of CNA associations to immune state were broad.

**Figure 2.**
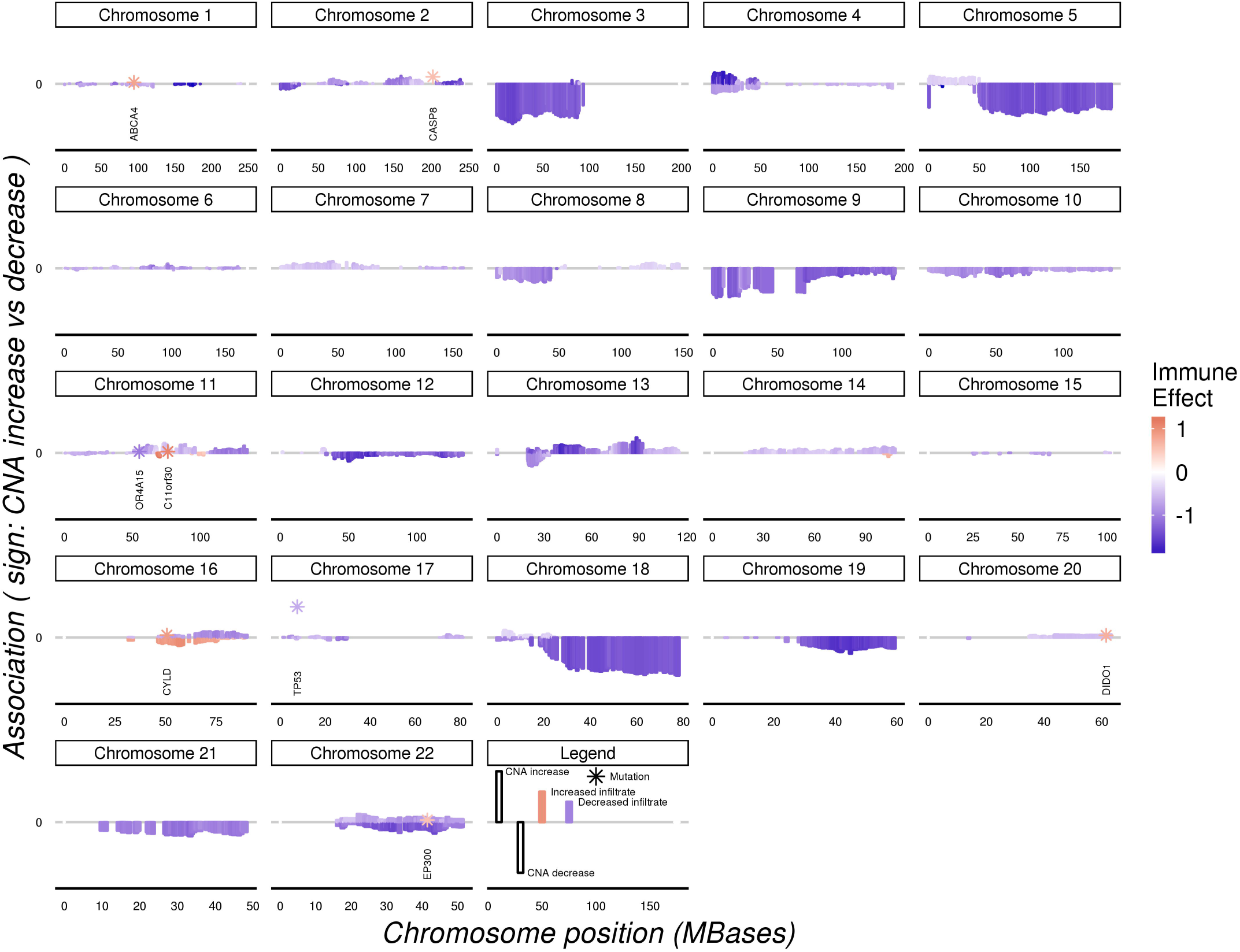
Landscape of association between tumor copy number changes, mutations, and CD8+ T cell estimates in TCGA head and neck cancer. Chromosomal location is shown on the horizontal axis with each point (mutation) or bar (CNA) representing the results for a locus. The length of the bars reflects the strength of the association signal; for CNAs, the sign indicates copy number gains (positive) or losses (negative). Mutation are indicated by stars and annotated with the HGNC gene name.

Among the well-studied drivers of oncogenesis we observed a strong link between copy number loss of the CDKN2A region of chromosome 9p and estimates of many immune cell types. CDKN2A is a tumor suppressor/negative regulator in the CDK4/Rb pathway, and is frequently lost and/or mutated in melanoma, pancreatic cancers, and other tumor types[25]. We observed a marked reduction in estimates of CD8+ T cells, Tregs, B cells, and the general T cell population linked to loss of the chromosome 9p region (Fig 3A-B). The associations between 9p loss and decrease in the estimated levels of infiltrate were observed in several tumor types, with the strongest associations seen in melanoma, pancreatic, and head/neck cancer cohorts (Table 2).

**Figure 3.**
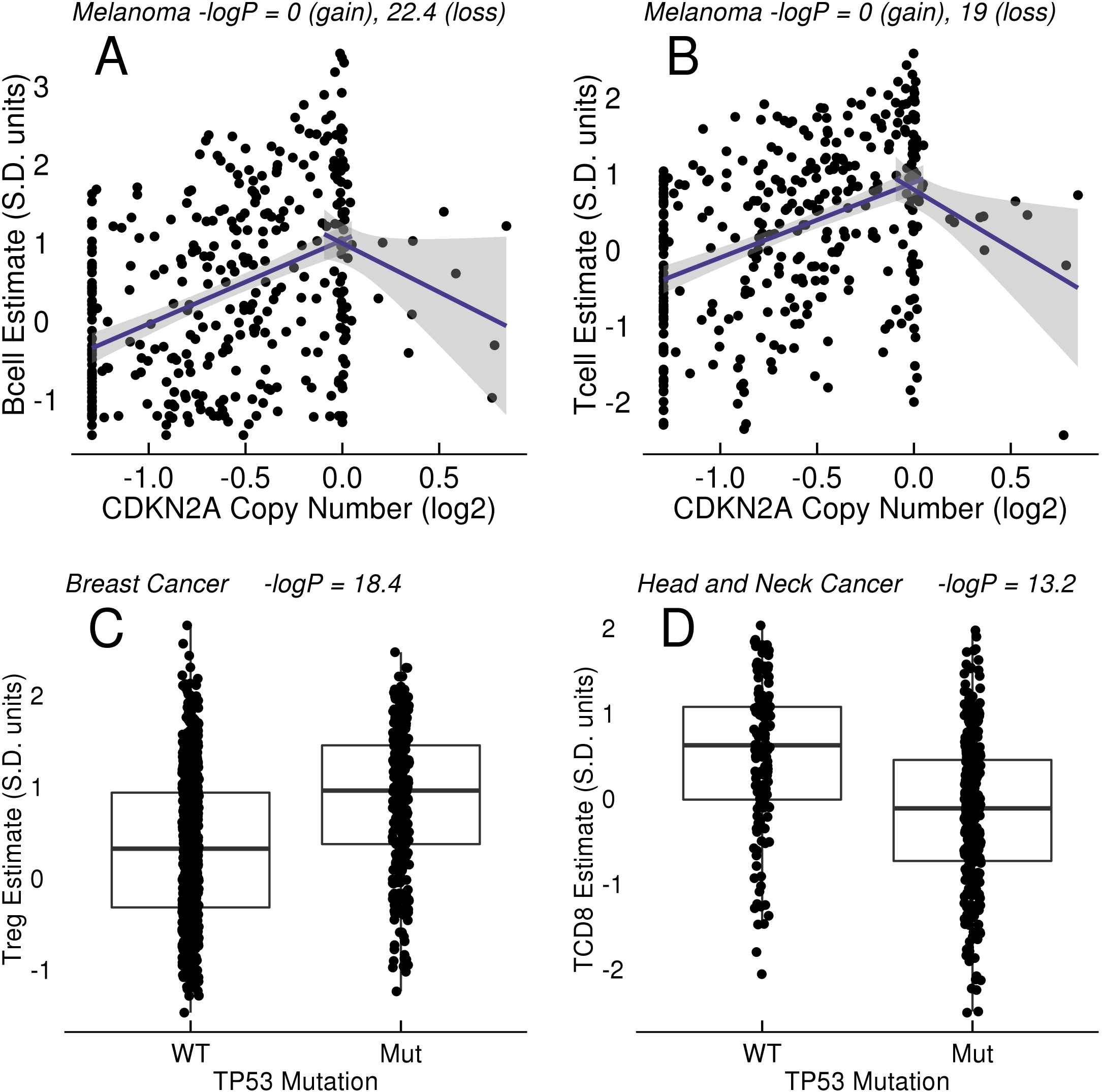
Association of CDKN2A CNA and TP53 mutation with immune estimates in tumors. (A-B) Relationship of CDKN2A copy number estimates to B and T cell estimates across TCGA melanoma. The horizontal scale is the log2 GISTIC CNA estimate (0 = diploid, -1.3 = homozygous loss). The signature scores are measured in units of standard deviation of the signature’s variation across TCGA tumors. Independent tests of association were performed for CNA > -0.1 and CNA < 0.1. The lines drawn are the linear regressions of the gain/loss CNA with the immune estimate, with shading to indicate the 95% confidence interval around the line’s slope (without model covariate adjustments or multiple test corrections). (C) Relationship of TP53 mutation to regulatory T cell (Treg) estimates across breast cancer. (D) Relationship of TP53 mutation to CD8+ Tcell estimates in head and neck cancer.

The region of association spans a large part of chromosome 9p and includes many loci in addition to CDKN2A (Fig 4). The genes in this region include a large cluster of genes encoding alpha-interferons and MTAP, a protein involved in adenosine metabolism that can also regulate STAT signalling[26]. The genes encoding PD-1 ligands CD274/PD-L1 and PDCD1LG2/PD-L2, as well as nearby Janus kinase JAK2, are sometimes contained within the region of 9p loss. The largest effect size on CD8+ T cell estimates were not at CDKN2A. We attempted to further dissect the region of association by analysis of significance, effect size, correlation of transcription with CNA, and concordance of these signals across multiple tumor types. For example, we examined chromosome 9 across melanoma, head/neck, and pancreatic cancer, using effect size rather than significance as a measure, and also studying a mean-based summary of the three indications (S1 Fig). The analysis did not result in discovery of a peak of association that would suggest a candidate mediator, but the multi-indication analysis may have limited the list of top candidates to a smaller region of 9p. Finally, amplification of PD-L1 (CD274) on chromosome 9p by neoplasms is well documented in Hodgkin lymphoma; one might hypothesize that, considering its known immuno-supressive role, amplification of the PD-L1 genomic region could be an active immuno-evasion mechanism in multiple tumor types[27]. We did not observe evidence for significant association of PD-L1 copy number gains with altered immune estimates in melanoma (P = 1), lung adenocarcinoma (P = 1), or any other cohort in the solid tumors studied.

**Figure 4.**
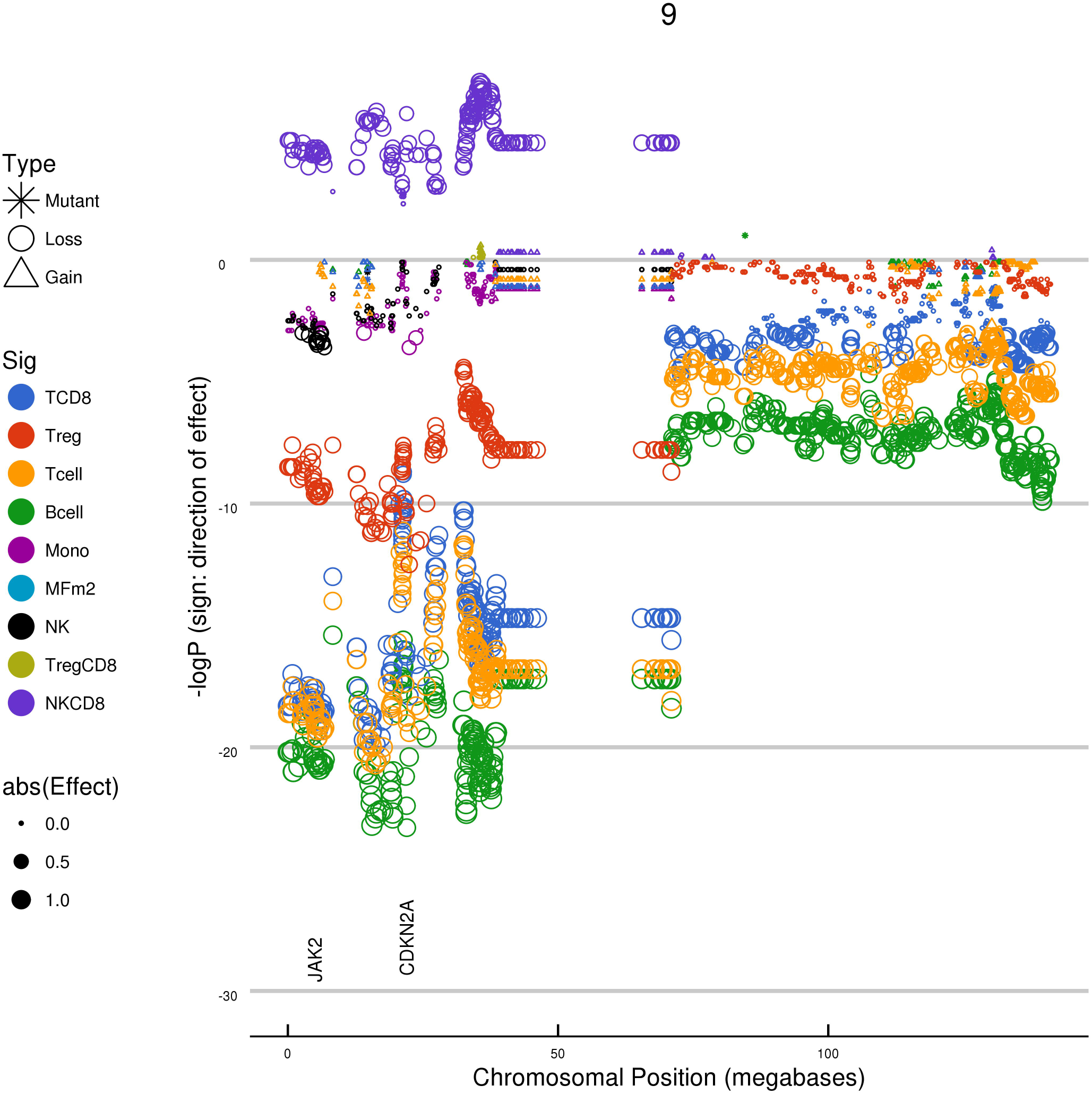
Relationship between chromosome 9 genetic changes and immune cell abundance estimates in TCGA melanoma. Chromosomal location is displayed on the horizontal axis, and effect size is displayed on the vertical axis. Each data point represents the results for a given locus, with significance (negative log(10) P value) indicated by the size of the data point. The negative log(10) of the multiplicity-corrected model P value is plotted on the vertical axes; negative values indicate a negative effect on the cellular estimate. A large region of chromosome 9p, when lost, is in association with the changes in cellular estimates for many immune cell types. The horizontal axis is the physical coordinate on chromosome 9 in units of 10^6^ bases. The vertical axis is the negative log(10) of the model P value, with negative numbers used to indicate associations that decrease the immune estimate being tested.

**Table 2.** Associations of loss of CDKN2A with estimates of immune cell types in tumors.

| Signature | Bladder | Breast | Head Neck | Kidney Clear | Lung Adeno | Melanoma | Pancreatic |
| --- | --- | --- | --- | --- | --- | --- | --- |
| Bcell | -0.38 (2.6) | -0.34 (2) | -0.62 (12) |  | -0.4 (2) | -0.88 (22.4) | -0.94 (7.2) |
| MFm2 | -0.35 (2.3) |  |  |  |  |  | -0.58 (1.5) |
| Mono |  |  | -0.35 (2.6) |  |  | -0.34 (1.6) | -0.59 (1.7) |
| NK |  |  | -0.38 (4.7) |  |  |  |  |
| NKCD8 |  |  |  |  |  | 0.5 (6) |  |
| TCD8 |  |  | -0.63 (12.5) |  |  | -0.76 (16.3) | -0.72 (3.3) |
| Tcell |  |  | -0.48 (7.9) |  |  | -0.76 (19) | -0.65 (3) |
| Treg |  |  | -0.58 (12.2) | 0.76 (5.9) |  | -0.58 (10.3) | -0.58 (1.9) |
| TregCD8 |  | 0.37 (3.1) |  | 0.74 (4.7) |  |  |  |

Values are reported as Effect Size with associated P values in parenthesis (-log10 P). The units of effect size are change in signature score (units of standard deviation) per unit of GISTIC (log2) CNA change.

TP53 mutation is the most common mutation in cancer, and loss of TP53 function leads to overall genetic instability and resistance to DNA damage-mediated apoptosis. We found greater immune cell abundance estimates in TP53 mutant breast cancer (Table 3, Fig 3C). However, TP53 mutation was associated with lower T, B, and NK cell abundance estimates in head and neck cancer (Fig 3D).

Several position-specific TP53 mutations were also found to be in association with estimates of T and NK cells. These mutations included R249S mutations, which have been linked to aflatoxin and hepatitis-associated liver cancer, and were associated with high estimates of NK cells in lung adenocarinoma[28]. Mutations in synaptonemal complex protein 2 (SYCP2), a protein involved in meiosis, were associated with lower estimates of Treg cells (-logP = 3, effect size -1.24) and the Treg/CD8 T cell ratio in head and neck cancer (Fig 5A.)[29].

**Figure 5.**
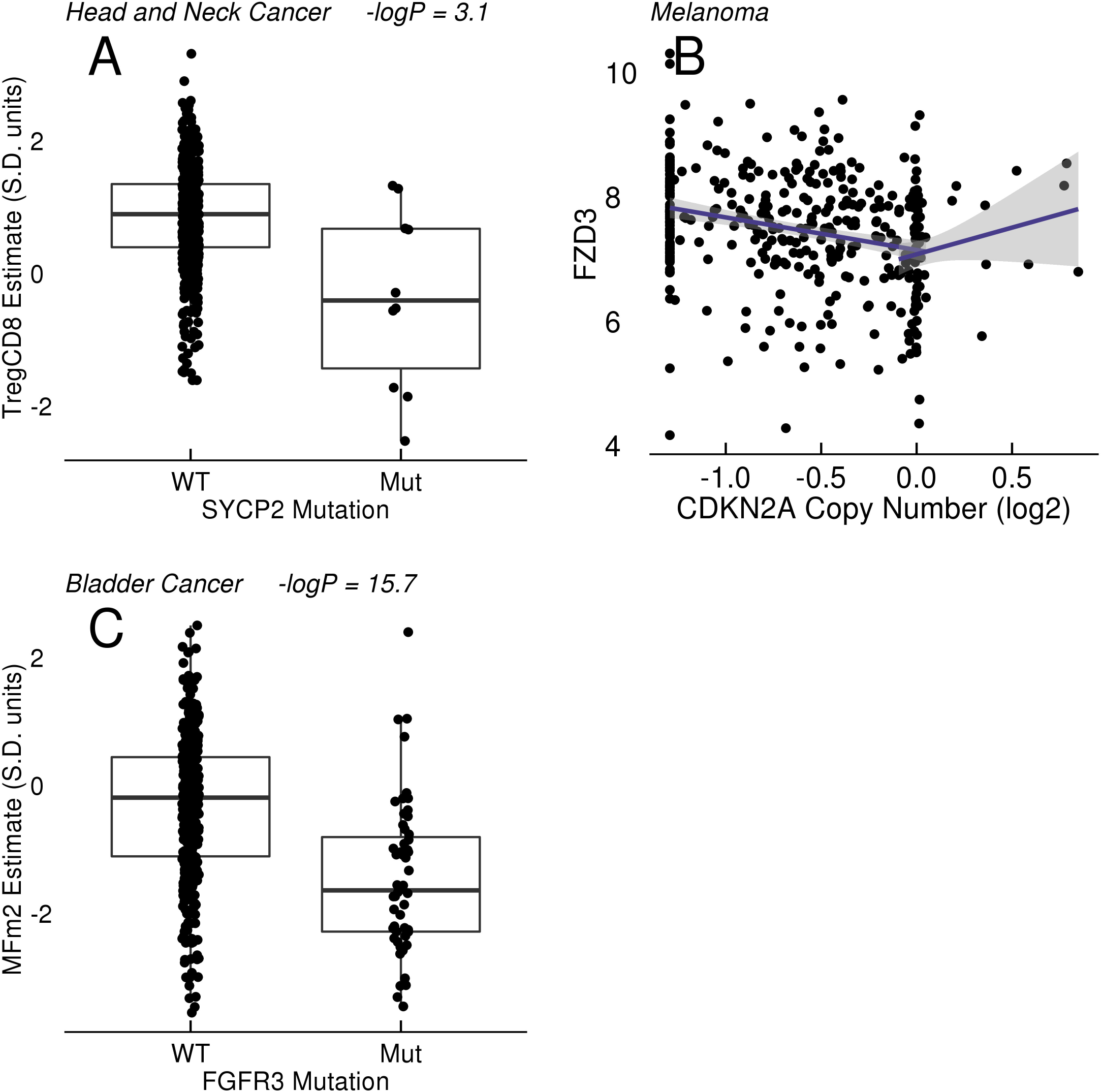
Association of SYCP2 and FGFR3 with immune estimates in tumors; correlation of chromosome 9p copy number (CDKN2A) with FZD3 RNA expression. A: Relationship between SYCP2 mutation and Treg - CD8 ratios in head and neck cancer. B: correlation of FZD3 (log2) RNA expression with CDKN2A copy number. C: Relationship between FGFR3 mutation and macrophage (MFm2) estimates in bladder cancer.

**Table 3.** Associations of TP53 mutations with estimates of immune cell levels across TCGA cohorts.

| Cohort | Sig | Variant | Association |
| --- | --- | --- | --- |
| Bladder Cancer | NK | TP53 G245S | 2.63 (4.2) |
| Breast (Basal) | NK | TP53 W91 stop | 2.91 (1.6) |
| Breast Cancer | Bcell | TP53 mutant | 0.29 (3.1) |
| Breast Cancer | NK | TP53 W91 stop | 3.24 (1.9) |
| Breast Cancer | NK | TP53 mutant | 0.26 (3.7) |
| Breast Cancer | TCD8 | TP53 mutant | 0.25 (1.7) |
| Breast Cancer | Tcell | TP53 mutant | 0.26 (2.4) |
| Breast Cancer | Treg | TP53 mutant | 0.51 (18.4) |
| Breast Cancer | TregCD8 | TP53 mutant | 0.26 (3.3) |
| Colon (MSS) | NK | TP53 S94 stop | 3.38 (6.2) |
| Colon (MSS) | NKCD8 | TP53 S94 stop | 3.02 (2) |
| Colon Cancer | NK | TP53 S94 stop | 3.17 (3.5) |
| Head and Neck Cancer | Bcell | TP53 mutant | -0.5 (6.1) |
| Head and Neck Cancer | NK | TP53 mutant | -0.48 (8.3) |
| Head and Neck Cancer | TCD8 | TP53 mutant | -0.66 (13.2) |
| Head and Neck Cancer | Tcell | TP53 mutant | -0.45 (5) |
| Head and Neck Cancer | TregCD8 | TP53 mutant | 0.41 (4.7) |
| Head and Neck Cancer | TregCD8 | TP53 N239D | -2.41 (2.5) |
| Lung Adenocarcinoma | NK | TP53 E224 splice | 4.46 (6.7) |
| Lung Adenocarcinoma | NK | TP53 R249S | 5.31 (7.5) |
| Lung Adenocarcinoma | NKCD8 | TP53 R249S | 4.33 (4.2) |
| Lung Adenocarcinoma | NKCD8 | TP53 E224 splice | 4.85 (6.9) |
| Lung Squamous Cell Carcinoma | NK | TP53 R158H | 3.14 (2.5) |
| Stomach Cancer | NK | TP53 I195N | 2.49 (2.8) |

Values are reported as Effect Size with associated P values in parenthesis ( $-\log_{10} P$ ). The units of effect size are change in signature score (units of standard deviation) for mutant versus wild type.

Among genes that may directly or indirectly modulate tumor immune state via interferon signaling or antigen presentation pathways, Zaretsky et al. describe loss-of-function mutations in Beta-2 microglobulin (B2M) and the JAK2 kinase as candidate mechanisms of acquired resistance to immunotherapy in the clinic[8]. B2M mutations were associated with higher estimates of NK cells in melanoma (effect size 1.3, -logP = 1.8) and lung adenocarcinoma (effect size 1.3, -logP 2 = 2). In micro-satellite stable (MSS) colon cancer we observed a similar positive association of JAK2 mutations with NK cell estimates (effect size 1.3, -logP = 1.7).

Recent reports suggest that activation of elements in the WNT/beta-catenin pathway lead to a suppressed immune micro-environment in melanoma models and possibly in human melanomas[13]. No significant association of beta-catenin (CTNNB1) mutations with estimates of immune levels was observed in our study, although there was a negative association between mutation and markers of fibroblast content in liver cancer (data not shown) Mutations of APC, a tumor suppressor in the WNT/beta-catenin pathway, were linked to lower levels of monocyte, macrophage, and CD8+ T cell estimates in colon adenocarcinoma and higher NK cell estimates in kidney papillary cell cancer (Table 4). Although one can observe negative correlations between MYC copy number gains and CD8 T cell estimates in melanoma (Pearson correlation: -0.18) and some other tumor types, the MYC copy number levels are also highly correlated with purity estimates (melanoma Pearson correlation: 0.33). We did not observe a significant association of MYC copy number gains with immune estimates in melanoma in our models that include purity estimates as a covariate, but did observe associations of small effect size in head/neck, stomach, and the Luminal B subtype (LumB) of breast cancer (Table 4). Finally, we noticed a strong correlation between WNT receptor Frizzled 3 (FZD3) expression and 9p loss in melanoma (Fig 5C.)

**Table 4.** Associations of APC mutation and MYC copy number gains with estimates of immune cell levels.

| Variant | Signature | Breast | Breast (LumB) | Colon | Head and Neck | Kidney Papillary Cell | Stomach |
| --- | --- | --- | --- | --- | --- | --- | --- |
| APC mutant | MFm2 |  |  | -0.75 (5) |  |  |  |
| APC mutant | Mono |  |  | -0.77 (5.2) |  |  |  |
| APC mutant | NK |  |  |  |  | 5.09 (9.4) |  |
| APC mutant | NKCD8 |  |  |  |  | 4.81 (7.2) |  |
| MYC gain | Bcell |  |  |  | -0.27 (1.9) |  | -0.22 (1.9) |
| MYC gain | MFm2 |  | 0.2 (2.2) |  |  |  |  |
| MYC gain | Mono |  | 0.19 (1.7) |  |  |  |  |
| MYC gain | TregCD8 | 0.1 (1.5) |  |  |  |  |  |

Values are reported as Effect Size with associated P values in parenthesis (-log10 P). The units of effect size are change in signature score (units of standard deviation) for mutant versus wild type in the case of mutations. For copy number changes the effect size is in S.D. units per change in GISTIC2 score.

Mutations in genes in the RAS/RAF pathway were associated with changes in immune estimates, but, despite the frequency of their occurence in different tumor histologies, we observed associations in limited tumor types. BRAF V600E mutations were associated with higher T cell and other immune cell estimates in thyroid cancer. (Table 5). Despite the high frequency of BRAF mutations in melanoma, we did not observe assocation of BRAF mutation with any immune signature there. Among the RAS family members, NRAS mutations were also linked to levels of T cells and Tregs in thyroid cancer. However, the direction of association was reversed from that of the BRAF mutations, with NRAS mutant tumors having lower estimates of CD8+ T, Treg, and pan-T cells.

**Table 5.** Association of RAS/RAF oncogene family member mutations with estimates of immune cell types across TCGA.

| Variant | Signature | Bladder | Colon | Testicular | Thyroid |
| --- | --- | --- | --- | --- | --- |
| BRAF mutant | MFm2 |  |  |  | 0.4 (2.7) |
| BRAF mutant | Mono |  |  |  | 0.45 (3.9) |
| BRAF mutant | NK |  | 0.58 (1.4) |  |  |
| BRAF mutant | NKCD8 |  |  |  | -0.38 (3.9) |
| BRAF mutant | Tcell |  |  |  | 0.48 (6) |
| BRAF mutant | Treg |  |  |  | 0.66 (14.9) |
| BRAF mutant | TregCD8 |  |  |  | 0.43 (5.9) |
| BRAF V600E | MFm2 |  |  |  | 0.45 (3.1) |
| BRAF V600E | Mono |  |  |  | 0.54 (5.2) |
| BRAF V600E | NKCD8 |  |  |  | -0.37 (3.1) |
| BRAF V600E | Tcell |  |  |  | 0.59 (10.1) |
| BRAF V600E | Treg |  |  |  | 0.75 (19.8) |
| BRAF V600E | TregCD8 |  |  |  | 0.49 (7.9) |
| KRAS G12V | MFm2 | -2.81 (2.8) |  |  |  |
| NRAS mutant | Bcell |  |  |  | -0.8 (3.7) |
| NRAS mutant | Tcell |  |  |  | -0.82 (4.7) |
| NRAS mutant | Treg |  |  |  | -0.74 (3.6) |
| NRAS Q61R | Bcell |  |  |  | -0.81 (2.2) |
| NRAS Q61R | NK |  |  | 1.64 (2.4) |  |
| NRAS Q61R | Tcell |  |  |  | -0.82 (2.8) |
| NRAS Q61R | Treg |  |  |  | -0.7 (2.2) |

Values are reported as Effect Size with associated P values in parenthesis (-log10 P). The units of effect size are change in signature score (units of standard deviation) for mutant versus wild type.

Mutations in the PIK3CA oncogene were associated with higher estimates of CD8+ T cells and NK cells, along with decreased Treg/CD8 ratios, across multiple tumor types (Table 6). We observed association with both gene-level mutation calls as well as position-specific mutations, many of which have been characterized as activating mutations in PIK3CA[30]. Mutations in FGFR3 were associated with decreased estimates of multiple immune infiltrate types in bladder cancer and increased NK cell estimates in head and neck cancer (Table 7 and Fig 5D). Many of the FGFR3 mutations introduce cysteines into the receptor that result in covalent dimerization and activation of the receptor[31] while others are reported to inhibit receptor internalization and enhance signalling[32].

**Table 6.** Relationship of PIK3CA oncogene mutations with estimates of immune cell types across TCGA.

| Cohort | Signature | Variant | Association |
| --- | --- | --- | --- |
| Breast Cancer | NK | PIK3CA E81K | 2.43 (2.4) |
| Cervical Cancer | NK | PIK3CA H1047R | 3.53 (3.3) |
| Colon (MSS) | NK | PIK3CA R88Q | 1.16 (1.7) |
| Stomach Cancer | TCD8 | PIK3CA mutant | 0.57 (2) |
| Testicular Cancer | NK | PIK3CA mutant | 1.9 (2.1) |

Values are reported as Effect Size with associated P values in parenthesis (-log<sub>10</sub> P). The units of effect size are change in signature score (units of standard deviation) for mutant versus wild type.

**Table 7.** Association of FGFR3 mutations with estimates of immune cell types across TCGA.

| Variant | Signature | Bladder | Head and Neck |
| --- | --- | --- | --- |
| FGFR3 mutant | Bcell | -0.71 (5.3) |  |
| FGFR3 mutant | MFm2 | -1.09 (15.7) |  |
| FGFR3 mutant | Mono | -1 (12.5) |  |
| FGFR3 mutant | Tcell | -0.51 (1.8) |  |
| FGFR3 mutant | Treg | -0.72 (6.9) |  |
| FGFR3 G382R | NK |  | 4.61 (5) |
| FGFR3 S249C | MFm2 | -0.86 (1.9) |  |

Values are reported as Effect Size with associated P values in parenthesis (-log<sub>10</sub> P). The units of effect size are change in signature score (units of standard deviation) for mutant versus wild type.

Finally, non-synonymous mutations in two distinct regulators of cell death were linked to altered immune state. Caspase 8 (CASP8) mutations were associated with higher T and NK cell estimates and lower Treg/CD8 ratios in head/neck cancer, and a specific Q156 nonsense mutation was associated with higher NK cells in breast cancer. (Fig 6A and Table 8). Death inducer-obliterator 1 (DIDO1) mutations were similarly associated with higher NK cell estimates (Fig 6B) in head/neck cancer. The pattern of mutations in across these genes was generally consistent with loss of function (data not shown), although some CASP8 mutations have been shown to increase nuclear factor kappa B signalling in tumor models[33].

**Figure 6.**
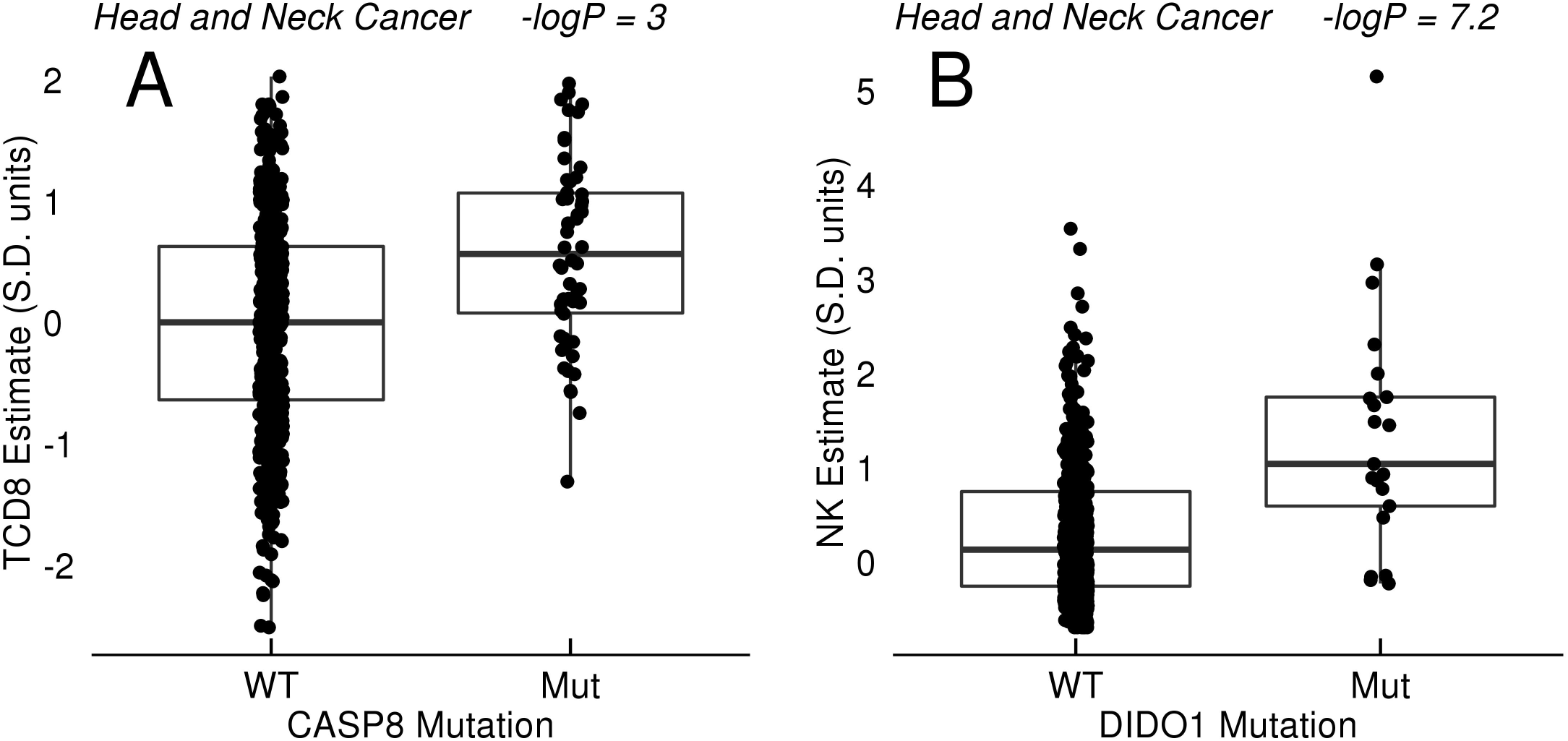
Mutations in cell death pathways. A: Relationship between CASP8 mutation with CD8+ T cell (TCD8) estimates in head and neck cancer. B: Relationship between DIDO1 mutation and NK estimates in head and neck cancer.

**Table 8.**
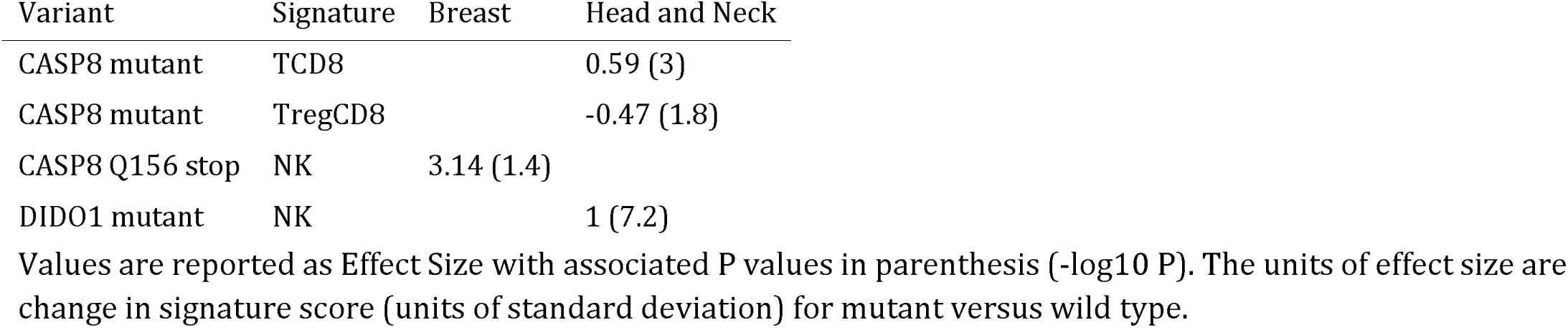
Association of mutations in cell death pathway genes CASP8 and DIDO1 with estimates of immune cell types across TCGA.

## Discussion

Investigating the association between between cancer genetics and tumor immune infiltrate requires quantitative estimates of intra-tumoral immune cell content. Of the various high-throughput data available in TCGA, RNA-seq data on transcription of immune markers is an obvious starting point, but not without challenges. Signature sets derived from RNA-seq of sorted immune cell populations from peripheral blood, such as FANTOM consortium studies, offer one way to estimate immune levels[21]. We were discouraged by our observation that only a fraction of markers derived from FANTOM data were well correlated with each other across tumors in TCGA. Markers derived from RNA-seq on sorted immune cells must demonstrate selectivity of expression not just versus other immune cells, but also across diverse tumor cells, tumor-associated stroma, and vasculature. A related challenge observed with immunological markers selected from the literature is that they are often derived from flow cytometry data; unfortunately we do not have the luxury of gating for a particular cell type with gross tumor RNA-seq data. Even a marker as canonical as CD4 is not a selective transcriptional marker for CD4+ T cells. Newman *et al*. recently reported a compelling support vector machine model that was trained with mixtures of immune and tumor cells and successfully predicts immune composition in tumors, but the published methods have not been trained for use on RNA-seq data, and the methods are not freely available[34].

We chose to re-derive immune signature marker sets directly from TCGA tumor data using a handful of sentinel markers (FOXP3, CD8A, CD19, etc.) for immune cells and utilizing mutual rank distance metrics to expand the local gene expression neighborhoods[23]. The use of mutual rank distance measures is one effective way to identify strict gene neighbors in the context of a large overall correlation structure of immune markers in tumors. The principle of this metric is similar to the widespread use of ‘reciprocal best hits’ in DNA and protein sequence analysis to define gene orthologs across species[35], but applied here to gene expression neighbors in RNA-seq data. Alternatively, one could have used partial correlation theoretical methods to derive similar sets of strict gene neighbors[36]. In our efforts to derive a signature for regulator T cells (Tregs), we discovered a tight association between FOXP3 expression and the expression of the chemokine receptor CCR8 across TCGA tumors. Survey of the the immunological genome database (http://immgen.org) and other databases of immune gene expession would not have suggested that CCR8 was a selective marker for Tregs. Plitas et al. have recently confirmed the selective expression of CCR8 on Treg cells from human breast tumors, and argue that anti-CCR8 therapies to target Tregs may be a promising approach to cancer immunotherapy[37].

We purposely kept these signature sets small (usually 2-3 genes/signature) and the mathematical model simple (usually a median of z-scored gene expression values). Larger set sizes and more complicated models could conceivably result in better predictors, but as signatures become more complex it also becomes more difficult to understand the nature of their failures. Because these sets were derived from correlation within tumors, rather than from isolated immune cell populations, they have demonstrated some level of resistance to interference from expression by the complex cellular environment of tumor and stroma. As a final conservative measure, we excluded tests of association of immune signatures with genetics for any given cohort when the median Pearson correlation within the marker set was less than 0.45. Work is ongoing to acquire a compendium of tumor RNA-seq data combined with flow-cytometry based quantitation of tumor immune content, which can serve as a gold standard for further assessment and validation of our immune cell type signatures.

Of the major genetic events that were in association with T cell levels in tumors, we found that loss of the chromosome 9p genomic region (driven by p16/CDKN2A) was among most significant. This result is in agreement with a report by Linsley *et al*. that demonstrated a link between interferon gene cluster loss (adjacent to CDKN2A) and decreased levels of several immune signatures in melanoma[38]. These effects were not reported by Rooney *et al*. as associated with their cytotoxic T cell signature, possibly because a strong effect is only observed in a handful of cohorts across TCGA[15]. In general, we see no reason to presume that the driver of a tumor CNA (in this case CDKN2A) is necessarily also the driver of the immune effect. The region of association we observed for 9p loss is present across the entire chromosome arm. Within it are the adenosine-modulating enzyme MTAP and the largest cluster of alpha interferons in the human genome. In addition, the PD-1 ligands PD-L1 and PD-L2, as well as the JAK/STAT member JAK2, are on 9p and sometimes in linkage with CDKN2A loss.

The reported trends for higher immunotherapy response rates in tumors with higher infiltrate suggests that our results might be used to guide therapeutic options, regardless of our understanding of cause and effect relationships[4]. One could infer that tumors harboring 9p alterations, in addition to having fewer T cells, would be less responsive to immunotherapy. Zaretsky *et al*. have recently reported a genetic study focused on patients that relapse during the course of pembrolizumab (anti - PD-1) therapy[8]. Although the study was limited to a few patients, they observed homozygous loss-of-function mutations in JAK2 in a relapsed patient, and *in vitro* studies demonstrated that cell lines lacking JAK2 were incapable of responding to gamma-interferon. Also, Gao et al. have studied mechanisms of resistance to anti-CTLA4 therapy in metastatic melanoma and concluded that copy number alterations containing interferons and interferon pathway genes, many on chromosome 9p, can predict response to therapy[39]. Thus, an accumulating body of evidence is now pointing to genetic disruptions of chromosome 9p playing a role in resistance to immuno-therapy.

Our study independently assessed the effects of copy number gains and losses. We reasoned that the biological driver of copy number gains and losses observed in any chromosomal region could often be distinct. This allowed an analysis of copy number gains of PD-L1(CD274) and PD-L2(PDCD1LG2) on chromosome 9p, despite the partial linkage with nearby CDKN2A loss that would have resulted in a spurious association in a combined analysis. Expression of PD-L1 in tumors is associated with response rates to anti - PDCD1 therapy[40]. Amplification of PD-L1 by neoplasms is well documented in Hodgkin lymphoma, and one might hypothesize that amplification of the PD-L1 genomic region could be an active immuno-evasion mechanism in multiple tumor types[27]. However, we observed no compelling evidence for association of PD-L1 amplification with any immune cell abundance estimate tested.

We observed several other very large chromosomal regions whose copy number estimates were associated with abundance estimates for many immune cell types. Most of these events, such as CNA on the long arm of chromosome 5, are copy number losses linked to decreased estimates of lymphocyte abundance (Fig 2). This common pattern of CNA loss leading to decreased immune estimates might suggest that immuno-editing could be taking place. Further work will be needed to confirm our attempted removal of the bias that tumor purity introduces to GISTIC estimates, a bias that would lead to aberrant association of all CNA with lower estimates of immune infiltrate.

One might have expected that the genetic instability of tumors with inactivated TP53 would be associated with higher immunogenicity of tumor and higher immune infiltrate. We did not observe a general correlation of TP53 mutations with immune cell estimates in tumors across TCGA cohorts, but associations were present in breast and head/neck cancers. TP53 mutations were associated with higher estimates of immune infiltrates in breast cancer. However, we found that in head and neck cancers the presence of TP53 mutations was associated with lower estimates of various immune infiltrates. As TP53 mutation has been shown to be inversely correlated with human papilloma virus infection (HPV) status in head and neck cancer, we believe that TP53 mutation status may be serving as an inverse marker of viral infection in this tumor type, with the HPV-infected tumors displaying a higher estimate of immune infiltrate[41]. In some cases we observed effects of specific TP53 mutations, such as the R249S mutation in lung adenocarcinoma, previously described in hepatitis and aflatoxin-associated liver cancer, that alter TP53 function in ways more subtle than simple loss of function and also suggests that some genetic-immune interactions could be related to environmental and/or viral insults[28].

Our study confirmed the report of Rooney *et al*. that caspase-8 mutations are associated with altered immune content estimates in some tumors. This is not an independent confirmation, as the data in our analyses both come from only modestly different releases of TCGA. However, in our study we observed a significant association only in head/neck and breast cancer[15]. CASP8 is one of the terminal elements in the cellular apoptosis pathway. CASP8 mutations were associated with higher CD8+ T and NK cell levels, in a mutational pattern suggesting loss of function. One possibility is that mutations of CASP8 are adaptations to an established immune response, providing resistance to T and NK cell-mediated cell killing. However, the switch from apoptotic to necroptotic cell death pathways can be associated with higher immunogenicity, especially when nuclear factor kappa B activity is present in the dying cells, which should lead to higher infiltrate[42]. It has also been reported that many CASP8 mutations, while loss of function in some aspects, result in increases in nuclear factor kappa B signaling and general tumor inflammatory state[33].

We also found that mutations in the cell death modulator DIDO1 (death obliterator-inducer 1) manifest a similar pattern in to CASP8 in head and neck cancer, with DIDO1 mutations associated with higher estimates of natural killer cells. Finally, we observe mutations in beta-2 microglobulin (B2M), a necessary component of antigen presentation, associated with high estimates of NK cells in melanoma, and JAK2 mutations with higher NK levels in colon cancer. These results are thematically in line with recent report of mutations in beta-2 microglobulin and JAK kinases as escape mechanisms in patients that have relapsed during pembrolizumab therapy[8] and consistent with the previous report of B2M association with immune state by Rooney *et al*.[15].

Although we saw some evidence of genetic changes in the WNT/beta-catenin pathway in association with immune estimates, the results are mixed. We did not observe compelling evidence for the association of any CTNNB1 (beta-catenin) mutation on immune estimates in our study. We did observe a link between the presence of APC mutations and decreases in estimates of myeloid and CD8+ T cells in colon adenocarcinoma, but increases in NK estimates in kidney papillary carcinoma were observed. Although amplification of MYC is inversely correlated with T cell estimates in some tumor types and Spranger et al. report a strong relationship between MYC expression and T cell estimates in melanoma[13], once we included estimates of tumor purity as a covariate in the analysis we did not observe association of MYC CNA with immune estimates in melanoma, and limited association elsewhere. We did, however, observe a strong correlation between FZD3 receptor expression and 9p loss in melanoma, suggesting that there may be a link between chromosome 9p loss and beta-catenin pathway status. To more thoroughly test WNT/beta-catenin hypotheses, multi-gene and multi-positional genetic signatures may need to be created to capture beta-catenin pathway activation via diverse genetic changes.

Although our study attempts to be comprehensive, it is limited in several aspects. A number of TCGA cohorts are still not large enough to expect sensitive identification of genetic-immune interactions, and even for large cohorts associations with rarer genetic changes will be too infrequent to sensitively measure. Although the rarer mutations and CNA might not be practically useful for patient stratification, they could still be a source of biologically rich information about the interactions of tumor genetics with immune state.

We attempted to adjust for two of the major possible biases in TCGA data in our models. First, one might expect that tumors with higher overall mutational burdens will have increased immune infiltrate, and this is observed in some tumor types[15,43]. We tried to control for this by using the observed total mutational burden of each sample as a covariate in our studies of mutation. It is possible that these corrections are over-conservative in cases where high mutational load and immune response leads to genetic adaptation by tumor, consistent with the observations of B2M mutations reported by Rooney *et al*.[15], where we observe a more limited strength of association in our model. Second, the GISTIC2 estimates of copy number alteration will be attenuated by the diploid nature of any non-tumor cell within the tumor. Left uncorrected, this bias can result in the prediction that any high level amplification or deletion will be immunosuppressive. We similarly chose to include an estimate of tumor purity as a covariate in our CNA model to try to adjust for this bias, despite the possibility the adjustment is overly conservative, as immune infiltrate is one component of tumor impurity. Finally, more intricate regression models than the ones described here could be devised that include factors such as clinical stage, sex, and more detailed tissue origin into the models.

It is important to highlight that we cannot infer cause and effect, in either direction, from any results in the current study. More conventional studies of human genetic effects on immune state usually are usually grounded in the assumption that the genetic variation is the cause of the immune change[44]. We cannot rely on that assumption in cancer. Although it is an intriguing hypothesis that the mutations in cell death pathways (CASP8, DIDO1) observed in head/neck cancer could be a tumor adaptation to ongoing immune activty, additional experiments will be necessary to establish this. There’s also the possibility that the genetic changes we observed are part of a rich history of tumor or tumor subtype evolution that ultimately and only indirectly lead to the immune change, i.e. we are identifying a latent tumor subtype. The observed association of TP53 mutations with lower immune estimates in head/neck cancer may be a reflection of the HPV origin of many of the TP53 wild-type tumors in the cohort. In thyroid cancer, the fact that activating BRAF and NRAS mutations are associated with immune changes in opposite directions, despite a common ability to activate the RAS/RAF/MEK/ERK pathway, may be another example where our analysis is identifying tumor subtypes rather than identifying cause-effect relationships. All of the above points mean that even for oncogene mutations linked to decreased infiltrate, like FGFR3 in bladder cancer, there is no assurance that inhibition of the oncogene’s signalling pathway will alter tumor immune dynamics. However, the FGFR3 result is one case where the cause and effect hypothesis is directly testable by study of FGFR3 inhibitors in cancer models.

An additional challenge, not uncommon in genome-wide association studies, is the sometimes very large chromosomal regions (CNA) found to be in association association with the phenotype (estimates of immune infiltrate). Although in some cases these CNA possess a peak when inspected for significance and/or effect size, in many cases the regions of association span millions of bases. The attempt to dissect the 9p region is one example: extensive study of the region across multiple tumor types failed to yield a clear candidate mediator of the effects (assuming a cause-effect relationship exists).

From comparison between all position-specific mutations across the TCGA cohorts and T cell estimates, there is little in our results to suggest that we have identified recurrent T cell neoepitopes from tumors that lead to consistently altered lymphocyte levels, although we purposely limited our study to the more frequent mutations across TCGA. We did note that the BRAF V600E mutation is distinctly associated with higher T cell estimates, but this is only observed in thyroid cancer, despite the prevalence of this mutation in melanoma and occurence in other tumor types.

## Conclusion

Our study of the relationship between tumor genetics and immune infiltrate is a starting point toward understanding how tumor evolution shapes the immune response and immune evasion, and possibly vice versa. We have identified some of the major genetic events linked to immune cell levels in tumors, some of which will likely influence response to immunotherapy. We developed a set of transcriptional markers for estimating relative levels of various immune cell types via a network correlation approach that was designed to resist interference from the heterogenetity of tumor tissue. We observed a strong relationship between copy number loss of a large region of chromosome 9p and decreased lymphocyte estimates in melanoma and several other tumor types. Although we could not identify specific loci on 9p responsible for the association, the recent reports of mutations in inteferon signalling pathways that lead to resistance to immunotherapy suggests that both the alpha-interferon cluster as well as JAK2 are candidate mediators, and that 9p alterations are relevant to response/resistance to immuno-therapy. We also noted associations of several cancer driver mutations in genes such as PIK3CA, BRAF, RAS family members, and FGFR3, with estimates of immune infiltrate, although these associations were often observed in limited tumor types. Finally, we observed that mutations in cell death pathway genes, CASP8 and DIDO1 were associated with higher immune infiltrate in head and neck cancers, and that several position-specific TP53 mutations possesed this phenotype. We think it is reasonable to consider that these mutations may be adaptations to ongoing immune activity in those tumors, and that examination of the combination therapy of agents that stimulate or modify cell death pathways with immune checkpoint blocking agents is warranted.

We also discovered some of the complexities in working with immune signatures and multi-modal tumor data. The large regions of association of many copy number alterations with immune estimates will require further studies to ascribe cause and effect on immune state to any particular loci. Biases in genomic data due to tumor purity and other possible factors can give strong association signals; when we removed some of these biases many signals (such as MYC amplification) were diminished or disappeared. Finally, the complexity of tumor composition and the highly correlated nature of immune infiltrate in tumors means that care must be taken to find markers as specific as possible for a given cell type. Efforts are underway to refine our marker sets as well as to acquire experimental data crucial to adding validation beyond our current criteria of mutual-rank correlation in tumors (manuscript in preparation).

Our genome-wide association analysis provides a baseline to compare against emerging hypotheses about tumor genetics and immunotherapy. As an example, in the case of WNT/beta-catenin pathway activation in melanoma, we were able to identify a relationship between what we view as a dominant effect of chromosome 9p loss with FZD3 receptor expression. We hope our work will help to guide future mechanistic studies on the influence of specific cancer pathways on immune state and response to immunotherapy, placing new hypotheses in the context of global tumor genetic - immune interactions.

## Methods

### TCGA data

Data from TCGA (TCGA Research Network: http://cancergenome.nih.gov/) was obtained from the University of California Santa Cruz Xena TCGA Pan-Cancer (PANCAN) repositories (http://xena.ucsc.edu/public-hubs/). Gene expression: UCSC Xena team, HiSeqV2_PANCAN, 2015-10-29. Values are log2(x+1) transformed RSEM gene-level expression estimates. CNA data: UCSC Xena team, TCGA_PANCAN_gistic2, 2015-10-26. Values are estimated using the GISTIC2 algorithm, the TCGA Firehose pipeline produced segmented CNV data, which was then mapped to genes to produce gene-level estimates. Gene-level Mutation data: UCSC Xena, TCGA_PANCAN_mutation_xena_gene, 2015-11-11. Genes are annotated as 0 (wt) or 1 (mutant) if they contain a non-silent mutation (nonsense, missense, frame-shift indels, splice site mutations, and stop codon read-throughs). Mutation were assigned a value of 0.5 if two different samples from the same tumor were analyzed and a single sample contained a non-silent mutation call. Position-level mutation data: UCSC Xena, TCGA_PANCAN_mutation_xena, 2015-11-11. Cancers of hematopoietic orgin (leukemias, lymphomas, gliomas, and thymomas) were excluded from the analysis.

Tumor subtype information for colon and breast cancers was taken from the corresponding phenotype annotation files from UCSC. Subtype classification for gastric cancer was obtained from the TCGA gastric cancer publication[41].

Analyses were performed in the R language for statistical computing [45]. The plyr package was used for data manipulation[46], ggplot2 was used for plotting[47], and limma was employed for model fitting and hypothesis testing[48,49]. The qgraph package was used for network plotting[50]. Knitr was used for manuscript generation from R Markdown[51–53].

Transcript co-regulation was measured using a 3-way mutual rank distance, an extension of the method described by Huttenhower *et al*. across all transcripts in the TCGA RNA data set, tumor only[23]. Each loci’s transcript expression values from TCGA PANCAN were first individually fitted to a standard distribution within cohort, then all cohorts were combined into a single data set. Pearson correlation was then used to rank all neighbors for each gene. All possible gene trios were evaluated for the minimum product of their six mutual ranks. Each gene-gene distance was expressed as the base 10 logarithm of this minimum score.

### Melanoma Single cell RNA-seq

Processed single cell RNA-seq data from Tirosh et al [20] were obtained from the Broad Single Cell Portal (https://portals.broadinstitute.org/single_cell).

### Signatures that estimate immune cell content in tumors

Signature scoring: Signature estimates were constructed as the median of z-scored (log2) expression values of each signature gene component except for the NK markers (see below).

TCD8 (CD8+ T cells): (CD8A, CD8B) Source: Mining of immune signatures in tumors using CD8A as sentinel marker. Reciprocal-Mutual-Rank methods were used to identify transcripts most intimately associated with sentinel markers. Caveats: CD8A is also expressed in a fraction of dendritic cells, some NK cells, and occasionally (rarely) in tumors.

Treg (Regulatory T Cells): (FOXP3, CCR8) Source: Mining of immune signatures in tumors using FOXP3 as sentinel marker. Reciprocal-Mutual-Rank methods were used to identify transcripts most intimately associated with sentinel markers. Caveats: Although CCR4 and CCR8 seem to be most predominantly co-expressed with FOXP3 in tumors, in sorted immune cells these receptors can also be seen in activated populations of CD4+ and CD8+ T cells.

Tcell (Pan T-Cell): (CD3D, CD3E, CD2) Mining of immune signatures in tumors using CD3 family members as sentinel markers. Reciprocal-Mutual-Rank methods were used to identify transcripts most intimately associated with CD3 epsilon (CD3E).

Bcell (B-cell): (CD19, CD79A, MS4A1) Source: Mining of immune signatures in tumors using CD19 as sentinel marker. Reciprocal-Mutual-Rank methods were used to identify transcripts most intimately associated with sentinel markers.

Mono (Monocyte lineage): (CD86, CSF1R, C3AR1) Source: Examination of correlation between antigen presenting cell-related genes across TCGA. Caveats: may not discriminate well between monocytes, macrophages, and other related members of the lineage.

M2mf (M2 Macrophage): (CD163, VSIG4, MS4A4A) Source: cross-referencing of Fantom/Hacohen/Rooney macrophage marker sets with mutual rank distance measures across TCGA[21]. The initial set was expanded with neighboring genes, cross-referenced with the literature and Mouse Immunological Genome Project (http://immgen.org) expression profiles to reduce to a small list of macrophage markers.

NK (Natural Killer cells): (KIR2DL1, KIR2DL3, KIR2DL4, KIR3DL1, KIR3DL2, KIR3DL3, KIR2DS4) Source: Mutual-rank correlation analysis of Natural Killer Group (NKG) and Killer-Cell Immmunoglobulin-Like Receptor (KIR) receptor families in TCGA tumor data revealed co-regulation of multiple members of the KIR family. However, any specific KIR gene was often observed to be at the lower limit of detection set by the TCGA RNA-seq pipeline. Compared to other cellular signatures, a larger collection of (KIR) markers was selected, a mean instead of median summarization was used to estimate NK cell content, and a small gaussian noise component was added (mean 0.16, standard deviation 0.08) to improve the normality of the NK signature score distribution.

TregCD8 and NKCD8 signatures were constructed by subtracting the TCD8 estimate from Treg estimate, or the TCD8 from the NK estimate, respectively.

### Analytical Models

Associations between gene mutations and immune signatures were estimated by the linear regression models of the form:

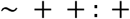

where ‘Total Mutation’ is the total number of observed mutations in the sample (log2 scaled), used as a covariate. ‘Mutation’ is considered a numeric variable in the model, with possible values of 0 (no mutation observed), 1 (mutation observed in TCGA sample), 0.5 (mutation observed in 1 of multiple TCGA samples available for that tumor).

Associations between gene copy number alterations and immune signature scores were estimated linear regression models of the form:

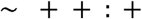

where copy number is a continuous-valued (log2) estimate obtained from GISTIC2 analysis of Affymetrix SNP 6 SNP/CNV array data, processed by TCGA[54].

Associations between copy number gains were performed with all data where GISTIC estimates were > -0.1 (log2 transformed GISTIC values). The associations with copy number losses were performed with data where GISTIC estimates were < 0.1.

Purity estimates were derived from data from the tumor purity meta-analysis published by Butte et al[24]. An RNA expression-based linear model was rederived from the Butte *et al*. purity estimates by fitting a 50 marker linear model of transcripts positively correlated with purity estimates, and applying the model to all samples in the study.

Mutation and CNA candidates: The top 3200 most frequently mutated genes across TCGA were used in the mutation analysis. All gene-level CNA were used in the analysis in order to allow finer mapping of association peaks. P values reported are corrected for multiple testing using p.adjust in R and the holm method[45] following the limma algorithm’s eBayes adjustment for multiple tests of immune signatures[48]. For mutation, the 3200 mutations in the analysis were used as the number of tests. For CNA, where groups of markers are often highly correlated, we single-linkage clustered all CNA data and identified 1023 separate groups that displayed pearson correlation less than 0.95 with each other. We used 1023 as the effective number of tests for the correction. In all of the models, we excluded tests of association of a particular immune signature with genetics for any given cohort when the median Pearson correlation within the marker set was less than 0.45.

## Acknowledgements

We thank the efforts of everyone involved in the creation of The Cancer Genome Atlas, especially the patients who made TCGA possible. We thank Mary Goldman and Jing Zhu from the University of California - Santa Cruz Xena team for their role in making TCGA data available to all, their expert help with interpretation of Xena TCGA data feeds, and responsiveness when we identified any concerns with the pan-cancer data. We also thank Itay Tirosh and co-authors of the single cell melanoma study for making the data readily available and helpful discussions. Support, discussions, and guidance of this research from Nils Lonberg, Alan Korman, Natalie Bezman, Ruth Lan, and Mark Selby at Bristol-Myers Squibb were invaluable.

## Supporting Information

**S1 Figure.**
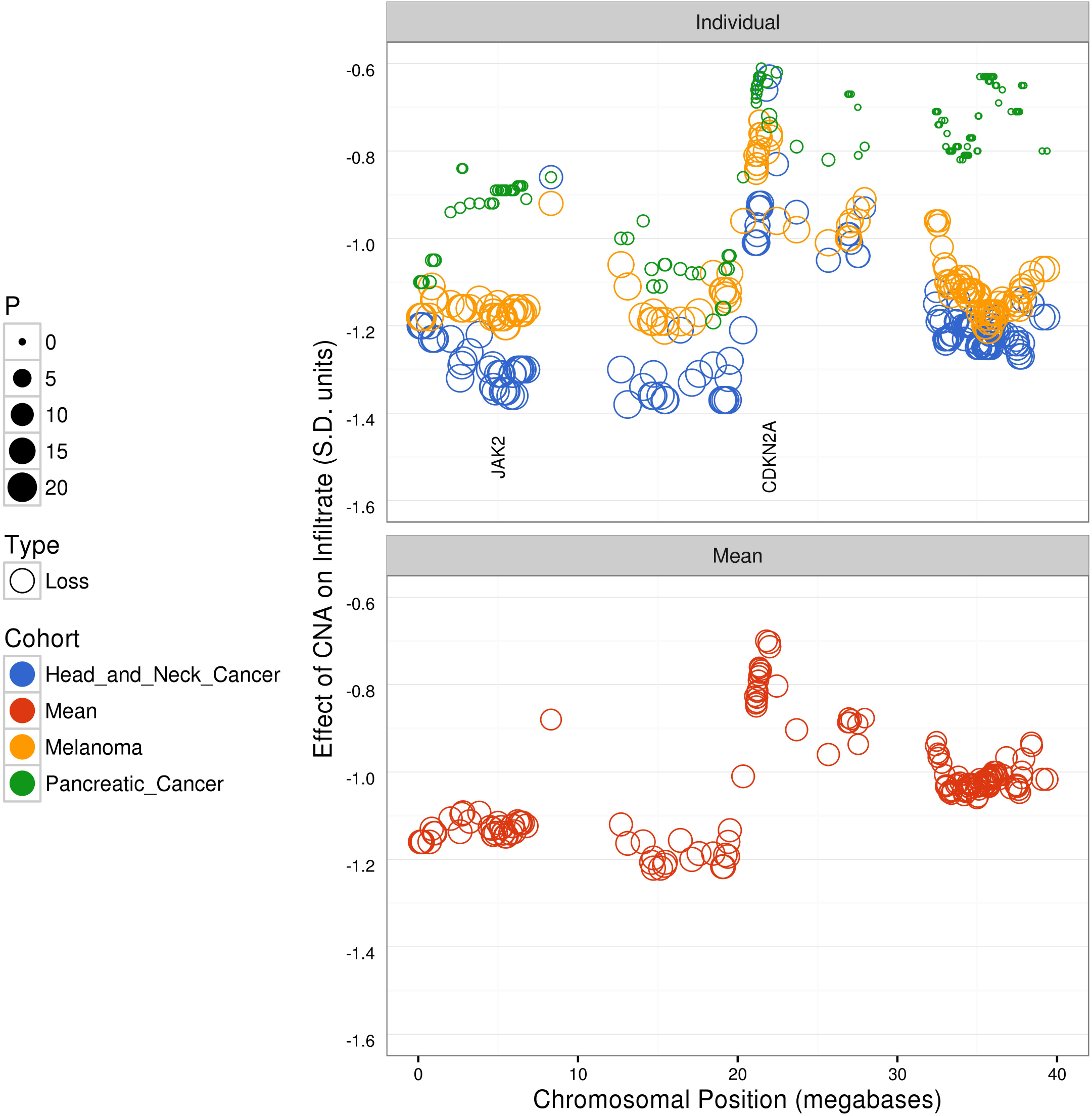
Alternative views of the relationship of chromosome 9 CNA to estimates of CD8+ T cell levels across melanoma, head/neck, and pancreatic cancer in TCGA. Chromosomal location is displayed on the horizontal axis, and the magnitude of effect of copy number change on CD8+ T cell estimates (rather than -logP, as in the main figures) is displayed on the vertical axis. Each data point represents the result for a given locus, with significance (negative log(10) of P value) indicated by size of the data point. The unit of effect size is the change in TCD8 signature score (units of standard deviation of signature score across all TCGA tumors) per (log2) unit of GISTIC copy number change. Individual panel: association between loss of a given chromosomal region and CD8+ T cell estimates in melanomaoma, pancreatic, head-neck cancer. Mean panel: a combined analysis across the three cohorts (mean effect size).

**Supplementary Data SData1.csv.gz** A compressed, comma-delimited file containing all the genetic association results contained in this study. Column key: “Cohort”, TCGA tumor type or sub-cohort tested; “Sig”, immune cell signature; “Type”, type of genetic change (CNA/cnv, mutation/mut, position specific mutation/fmut); “Variant”, mutation or CNA tested; “Chr”, chromosome of variant; “Pos”, position of variant(megabases); “minP”, minimum uncorrected P value from any tumor type among contrasts tested in model; “P”, corrected -log(10) P value; “Effect”, effect size; “Dir”, direction of effect size; “P.orig”, P value corrected for multiple tests of signatures by limma but not for multiple tests of genetic changes.

A snapshot of the UCSC Xena team’s November 2015 release of TCGA data used in this work, along with the R and R markdown code used to perform the analysis and generate this manuscript, is available at http://fiveprime.org/GENIO.

